# Mutational cascade of SARS-CoV-2 leading to evolution and emergence of omicron variant

**DOI:** 10.1101/2021.12.06.471389

**Authors:** Kanika Bansal, Sanjeet Kumar

## Abstract

**Background:** Emergence of new variant of SARS-CoV-2, namely omicron, has posed a global concern because of its high rate of transmissibility and mutations in its genome. Researchers worldwide are trying to understand the evolution and emergence of such variants to understand the mutational cascade events.

**Methods:** We have considered all omicron genomes (n = 302 genomes) available till 2^nd^ December 2021 in the public repository of GISAID along with representatives of variants of concern (VOC), i.e., alpha, beta, gamma, delta, and omicron; variant of interest (VOI) mu and lambda; and variant under monitoring (VUM). Whole genome-based phylogeny and mutational analysis were performed to understand the evolution of SARS CoV-2 leading to emergence of omicron variant.

**Results:** Whole genome-based phylogeny depicted two phylogroups (PG-I and PG-II) forming variant specific clades except for gamma and VUM GH. Mutational analysis detected 18,261 mutations in the omicron variant, majority of which were non-synonymous mutations in spike (A67, T547K, D614G, H655Y, N679K, P681H, D796Y, N856K, Q954H), followed by RNA dependent RNA polymerase (rdrp) (A1892T, I189V, P314L, K38R, T492I, V57V), ORF6 (M19M) and nucleocapsid protein (RG203KR).

**Conclusion:** Delta and omicron have evolutionary diverged into distinct phylogroups and do not share a common ancestry. While, omicron shares common ancestry with VOI lambda and its evolution is mainly derived by the non-synonymous mutations.

## Introduction

Currently, the world is witnessing a resurgence of COVID-19 cases due to the new omicron variant belonging to B.1.1.529. Omicron was first reported in South Africa on 24^th^ November 2021 from the specimen collected on 9th November 2021(https://www.who.int/publications/m/item/enhancing-readiness-for-omicron-(b.1.1.529)-technical-brief-and-priority-actions-for-member-states). On 26^th^ November 2021, World Health Organisation (WHO) assigned omicron to the ‘variant of concern’ (VOC) category due to its ability to poses a higher risk of reinfection as compared to previously reported variants (https://www.who.int/news/item/26-11-2021-classification-of-omicron-(b.1.1.529)-sars-cov-2-variant-of-concern; https://www.who.int/news/item/28-11-2021-update-on-omicron). According to the 1^st^ December 2021 update, omicron is reported in at least 23 countries from five out of six WHO regions, with most cases in Africa and Europe (https://www.cnbc.com/2021/12/01/who-says-omicron-has-been-found-in-23-countries-across-the-world.html).

There is a lot of uncertainty surrounding the omicron variant. For its risk assessment, scientists and researchers are investigating the intensity of its spread, extent of its infection, effectiveness of detection methods, therapeutics, and vaccine efficacy (Knoll & Wonodi, 2021; Lipsitch & Dean, 2020; Pegu et al., 2021). The onset of omicron is reported with mild diseases suggests its low or mild severity than its previous counterparts like delta (Ewen Callaway, 2021; E. Callaway & Ledford, 2021). It is known to have a very high mutation rate with more than 30 mutational changes in its spike protein (Ewen Callaway, 2021) (https://www.who.int/publications/m/item/enhancing-readiness-for-omicron-(b.1.1.529)-technical-brief-and-priority-actions-for-member-states)

Globally, high risk of reinfection with omicron variant, low vaccine and testing coverage are ideal for mutations resulting in the emergence of new variants of SARS-CoV-2 (Pulliam et al., 2021). Since COVID-19 inception, researchers have been trying to investigate its origin and evolution (Bansal, Kumar, & Patil, 2021; Singh & Soojin, 2021; Tang et al., 2020). We are currently witnessing a global molecular arms race between SARS-CoV-2 and its preventive therapeutics based on diverse regimes such as DNA, RNA, protein or inactivated whole-virion, etc. (Andreadakis et al., 2020; Corey, Mascola, Fauci, & Collins, 2020; Sharma, Sultan, Ding, & Triggle, 2020). This global crisis can be addressed by a very rapid immunization program worldwide. Moreover, the real-time monitoring of evolutionary cascade of SARS-CoV-2 leading to lethal variants is utmost. Earlier investigation of several VOC and VOI suggests some of the crucial mutations for viral survival and high infectivity in humans (Boehm et al., 2021; Kumar & Bansal, 2021; Schmidt et al., 2021). However, mutations giving rise to omicron and intra-omicron genomic diversity are not yet analyzed at a population level.

In the present study, we aim to look for the mutational profile of deadly and under-monitoring variants reported till now to understand the emergence of a heavily mutated variant named omicron. Interestingly, whole genome-based phylogeny suggests two major phylogroups PG-I and PG-II. Further, mutational analysis depicted the key role of non-synonymous mutations in the evolution of lethal variants. Such genomic insights into the nature of mutational cascade are the need of the hour.

## Results

### Phylogenomics suggests common ancestry of omicron and lambda variants

Whole genome-based phylogeny (n = 478 genomes) representing VOC (alpha, beta, gamma, delta, and omicron), VOI (mu and lambda), VUM depicts two major phylogroups PG-I and PG-II (figure 1, supplementary figure 1 and supplementary table 1). Here, the reference strain of SARS-CoV-2 (Wuhan-Hu-1, NC_045512.2) is taken as an outgroup. PG-I has VOC: gamma, beta, and delta; VOI: mu and VUM: GH. PG-II includes VOC: alpha, omicron and VOI: lambda. However, seven strains of VOC gamma are in PG-II, two of which are basal to PG-II forming its outgroup (EPI_ISL_3218258, EPI_ISL_2454057), and five (EPI_ISL_2220217, EPI_ISL_2216321, EPI_ISL_2223074, EPI_ISL_2224081, and EPI_ISL_2224090) are more related to alpha variant. EPI_ISL_3160245 one of the VUM strain is distant from the main clade of VUM.

**Figure 1:**
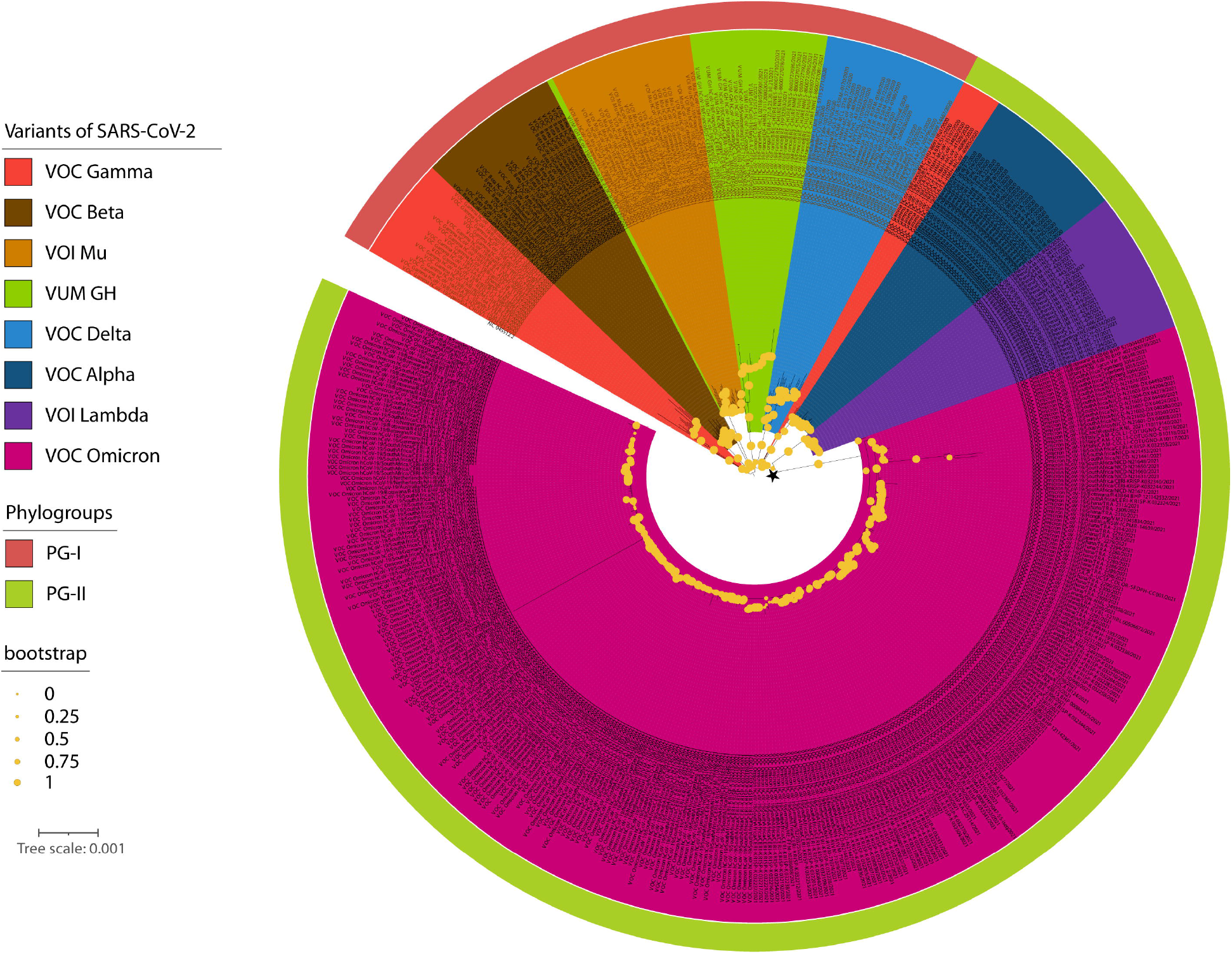
Maximum likelihood whole genome-based phylogeny of SARS-CoV-2 VOCs, VOIs and VUMs. Here, phylogroups (PG-I and PG-II) and clades (alpha, beta, gamma, delta, omicron, mu etc.) are marked with respective colors as indicated. Bootstrap values are represented by the radius of circle at the nodes. Common ancestry of omicron and lambda is marked by black star.

Interestingly, two deadly VOCs, delta and omicron, belong to different phylogroups. Phylogeny depicted that omicron shares a common ancestry with VOI lambda represented by a black asterisk in figure 1. Interestingly, three isolated from Italy (EPI_ISL_6854346, EPI_ISL_6854347, and EPI_ISL_6854348) form a diversified sub-lineage among the omicron population. Additionally, EPI_ISL_6886594 from Germany is a diversified omicron strain.

### Very high non-synonymous mutations give rise to omicron

To further understand the evolution and emergence of omicron, we have performed a mutational analysis with respect to the reference genome of SARS-CoV-2 (NC_045512.2) (figure 2). Total mutations detected in the dataset were 24,189, and omicron genomes constituted 18,261 (supplementary table 2). Interestingly, >97% (n = 17,703 mutations) of the mutations in omicron were in the coding region, and remaining 558 were detected in the extragenic region of the genome. Amongst the coding gene mutations, 2,965 were indels while 14,738 were SNPs constituting non-synonymous (n = 11,995 mutations) and synonymous mutations (n = 2,743 mutations).

**Figure 2:**
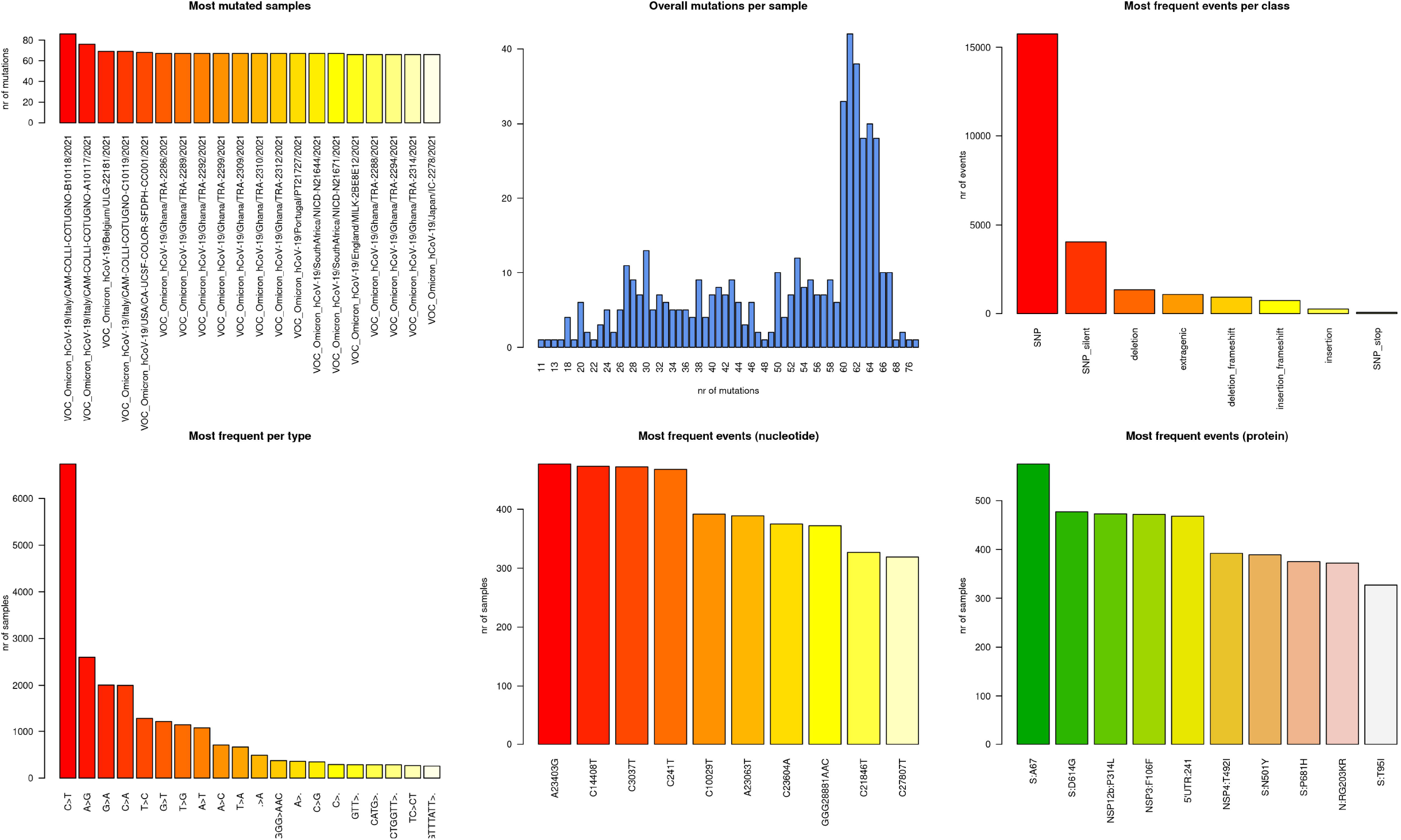
Mutational analysis of omicron. Six panel image displays the most mutated samples, overall mutations per samples, most frequent events per class of mutation category, changes of nucleotide per type, nucleotide wise most frequent events and protein level most frequent events for the genomes used in the study.

Interestingly, mutational events are highly skewed towards the spike protein, which constitutes ~60% (n = 10,658) of the total mutations in the coding genomic region (n = 17,703) (figure 3). The majority of spike protein mutations encompass A67, T547K, D614G, H655Y, N679K, P681H, D796Y, N856K, Q954H, which are reported in all the omicron genomes analyzed (table 3). Count of mutations in the spike was followed by RNA dependent RNA polymerase (rdrp) (n = 4,142) constituting A1892T, I189V, P314L, K38R, T492I, V57V in all omicron genomes analyzed (figure 3 and table 3). Remaining 2903 mutations were detected in rest of the coding genomic region (table 2, 3, and supplementary table 1), where M19M in ORF6, and RG203KR in nucleocapsid protein are amongst the most prevalent mutations in omicron (figure 3).

**Figure 3:**
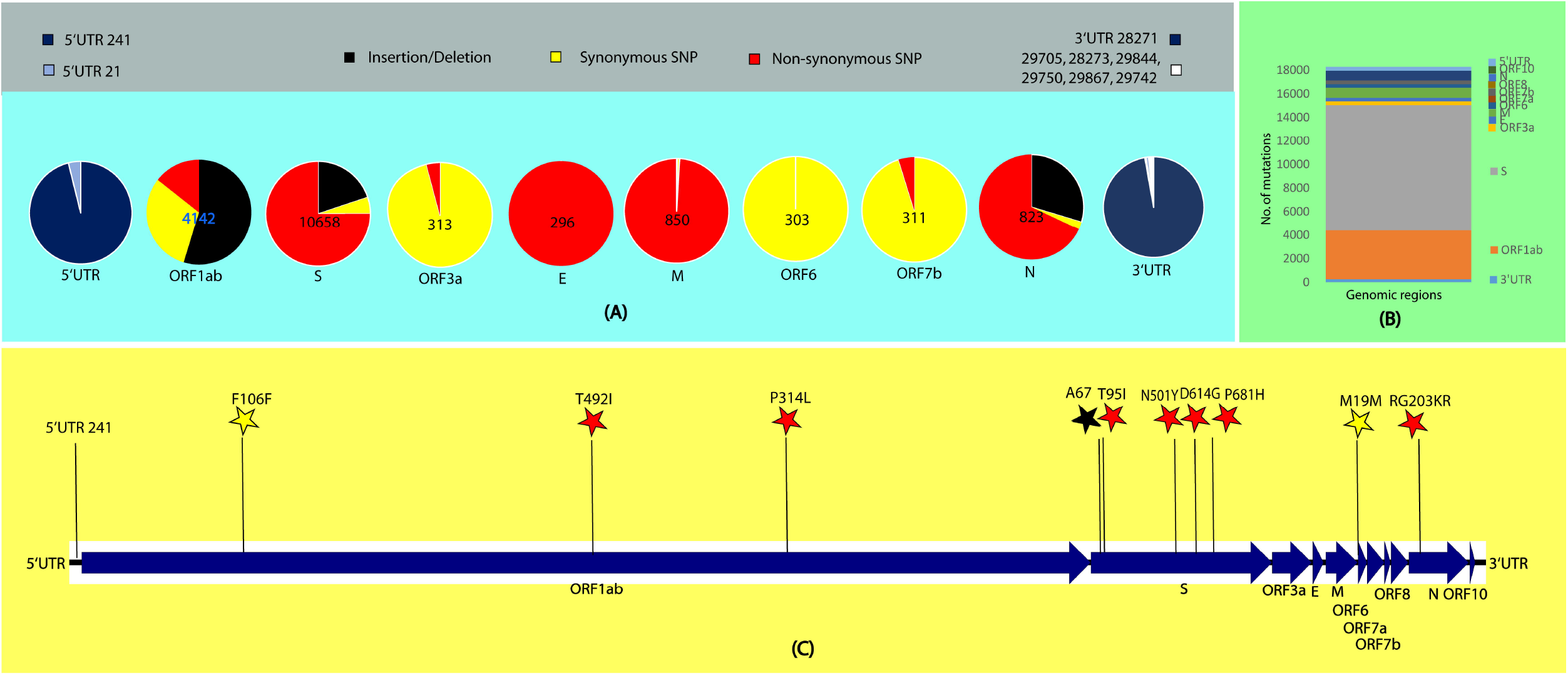
Mutational analysis of omicron. **(A)** Number of mutations in the coding region is in the centre of the pie-chart representing indels (black), synonymous (yellow) and non-synonymous (red) SNPs. Type and number of mutations in the extergenic region is represented by pie charts blue, light blue and white as represented in the color legends. **(B)** Bar graph representing number of mutations in the genomic region of SARS-CoV-2. **(C)** Some of the top mutations (pl. refer table 3 for all top mutations in omicron) among the omicron variant are represented by stars of black: indels, yellow: synonymous and red: non-synonymous mutations.

**Table 1:**
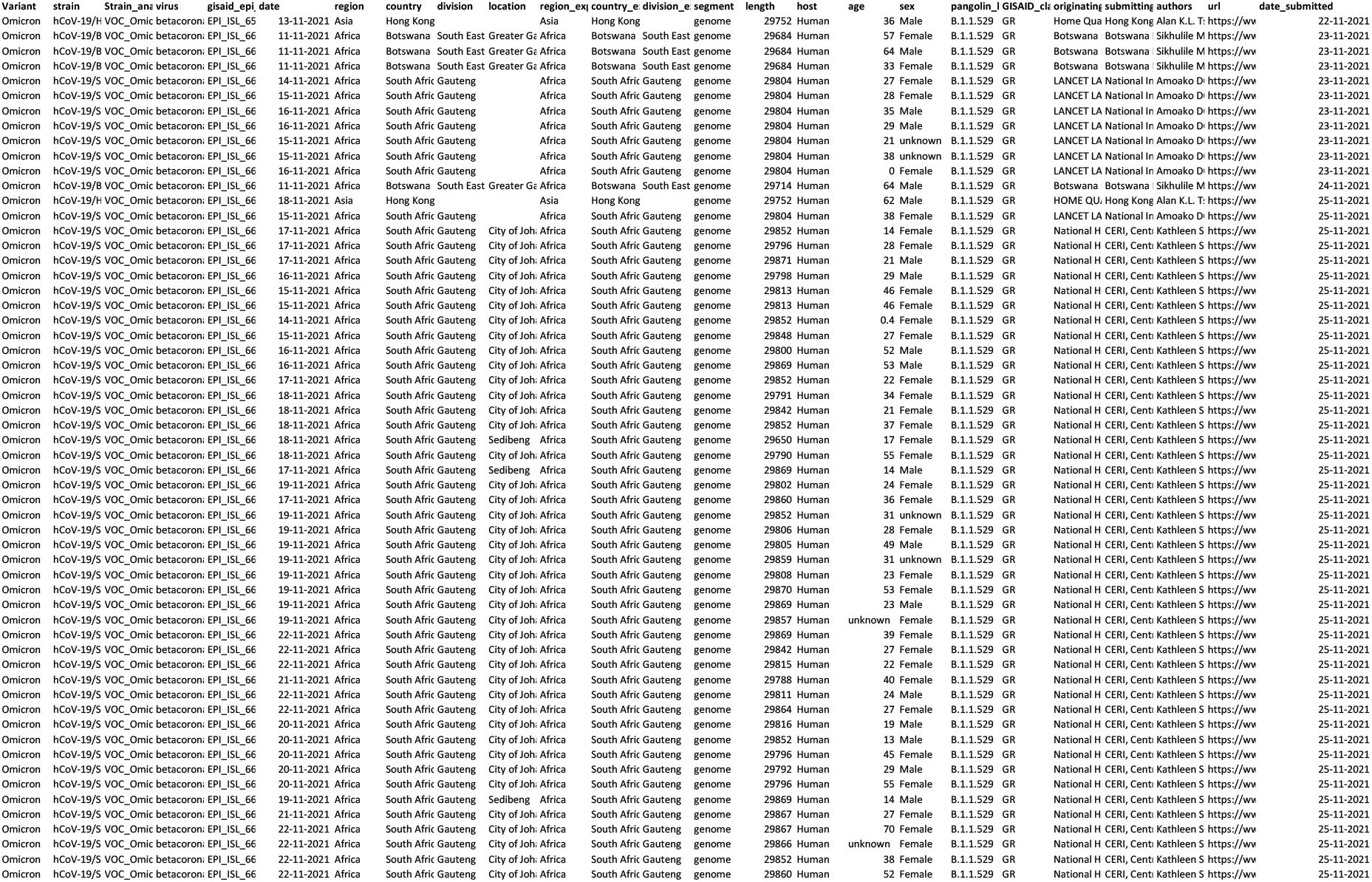
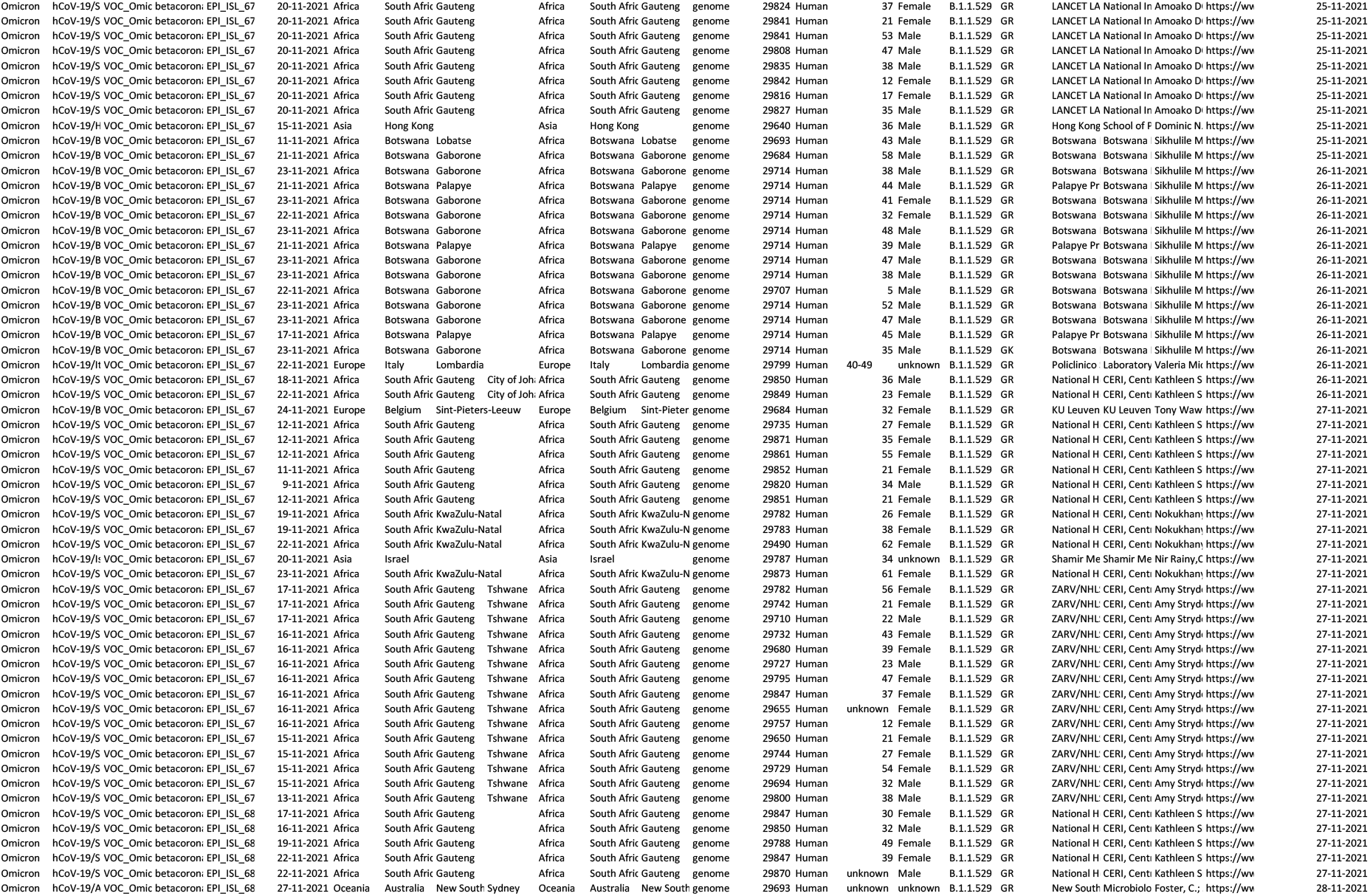
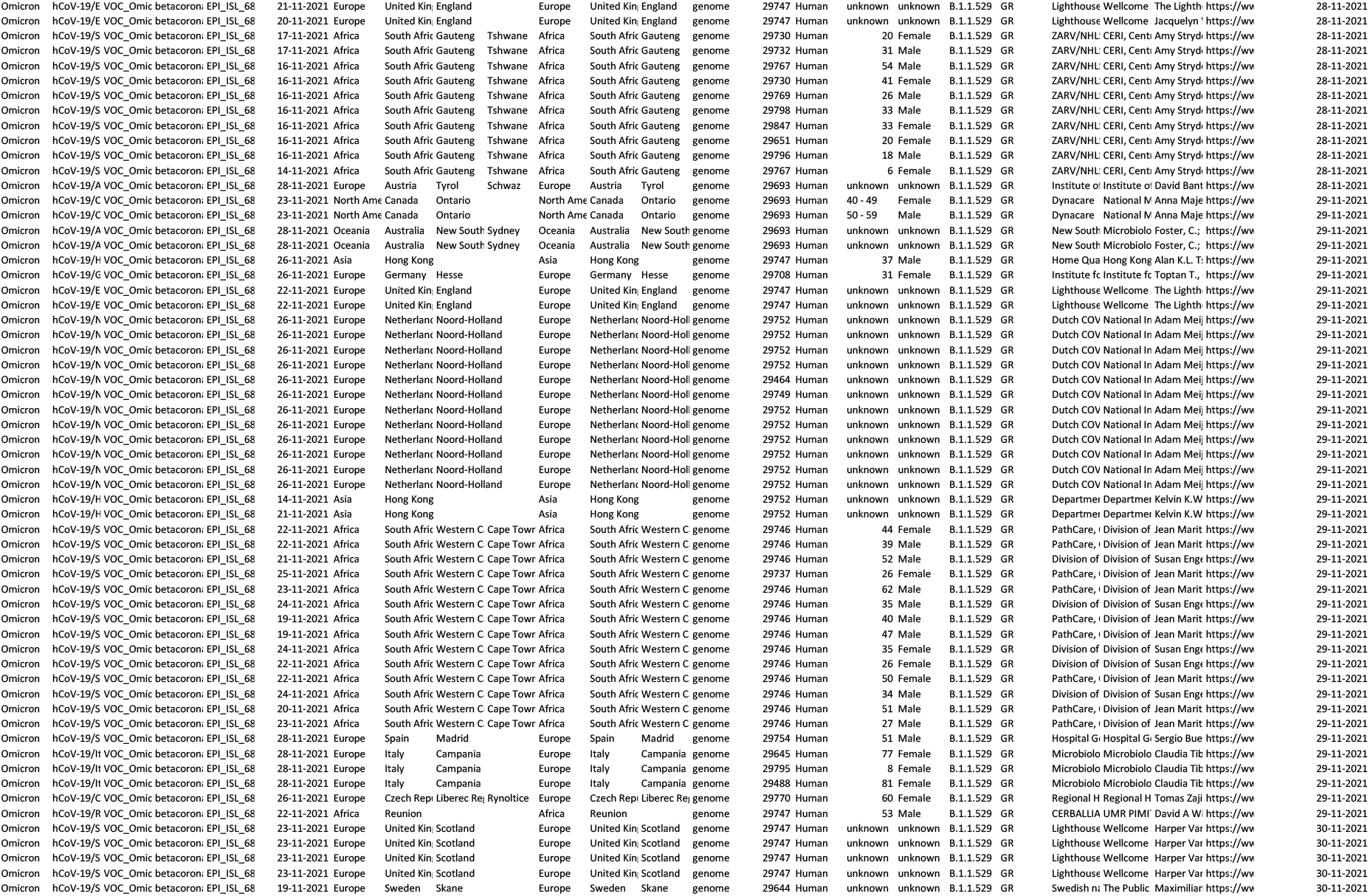
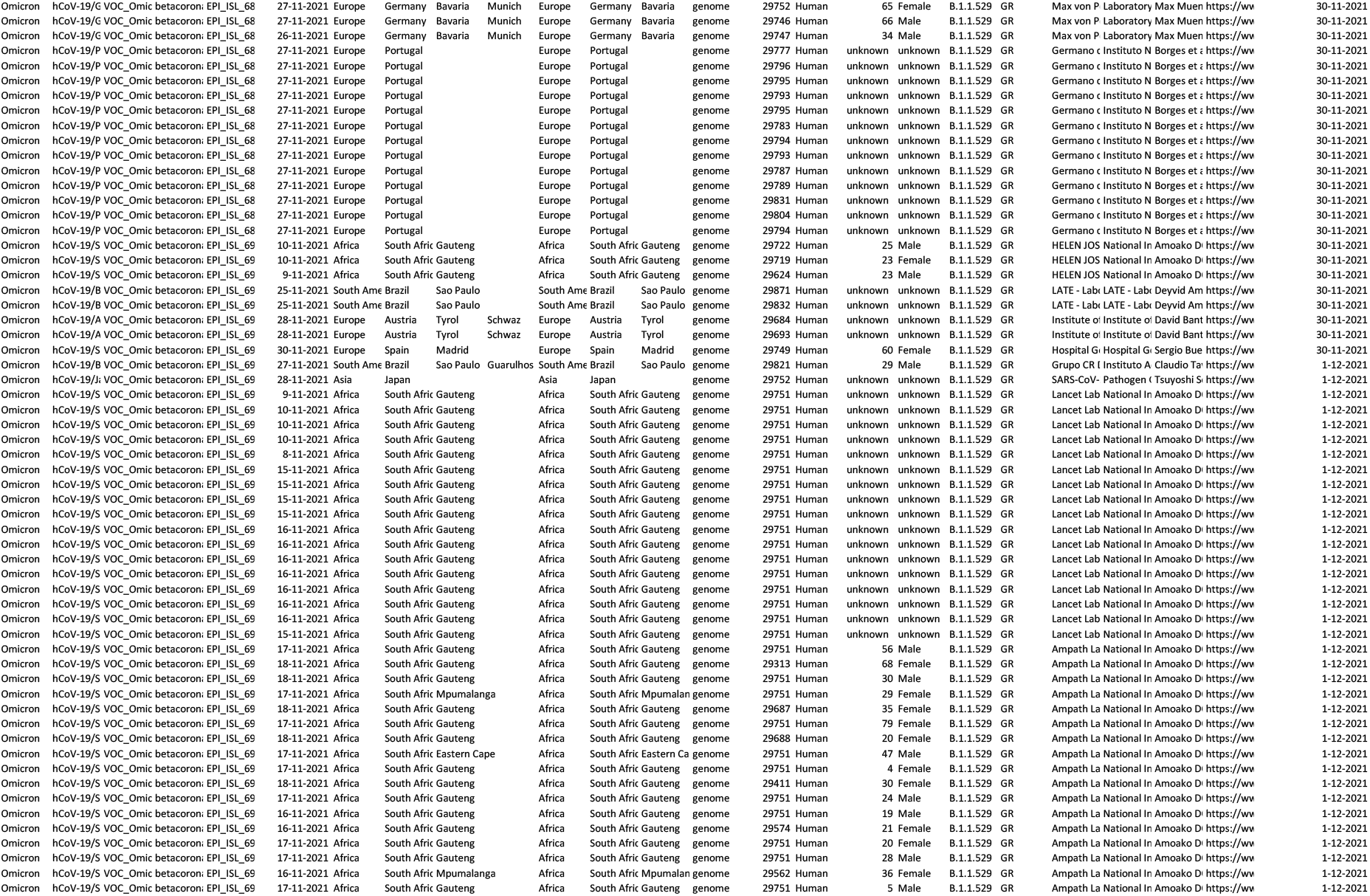
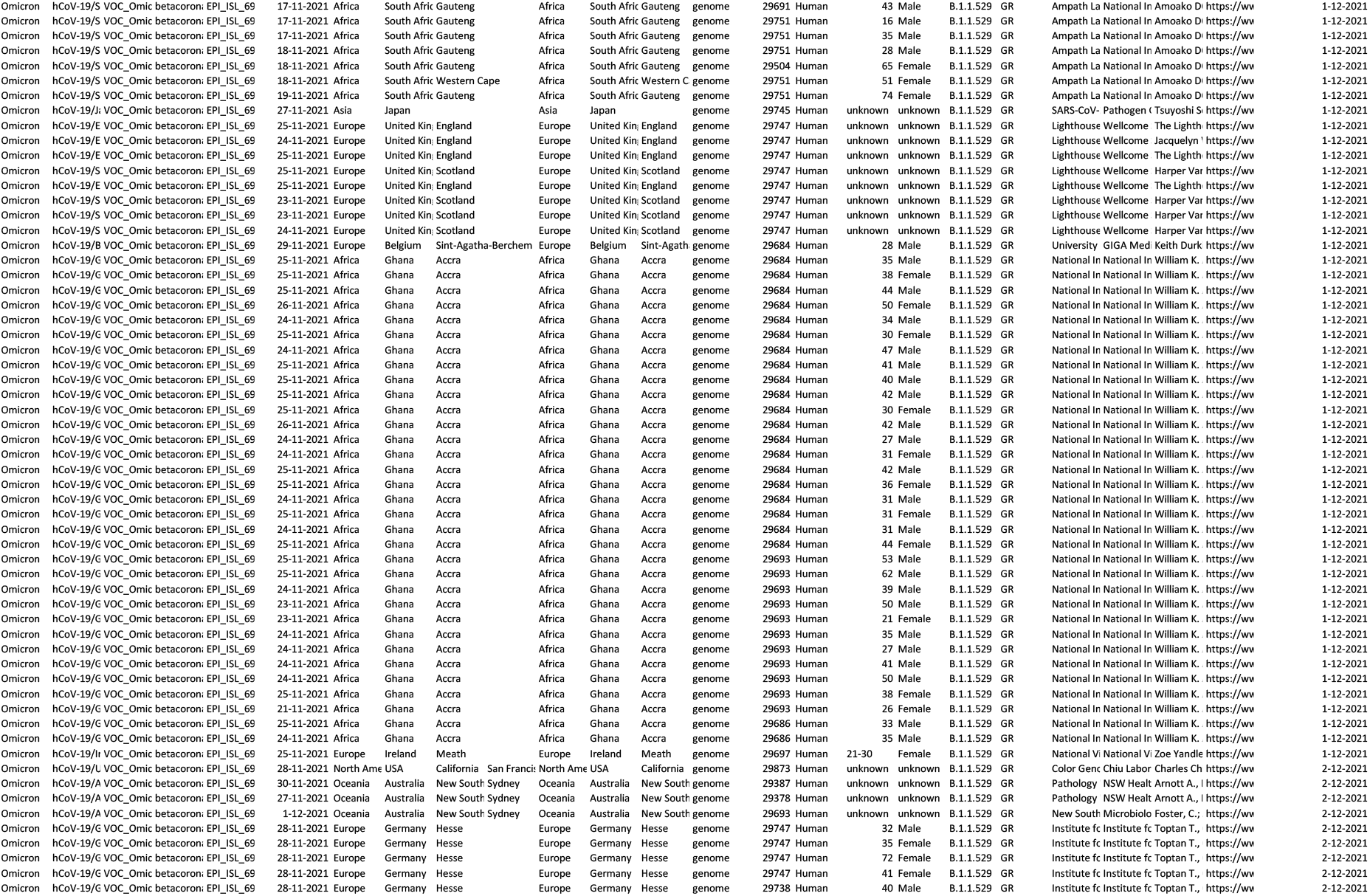
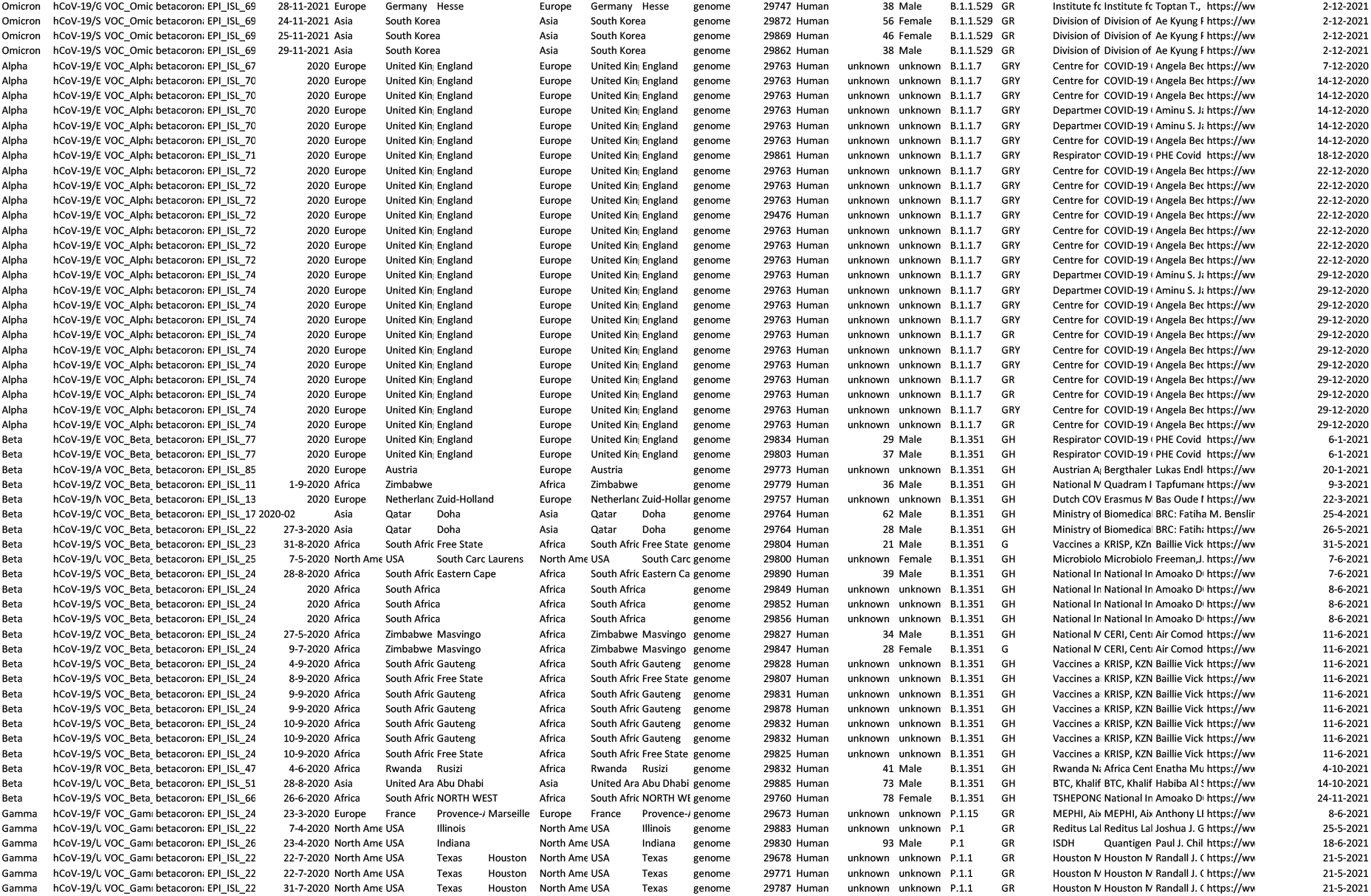
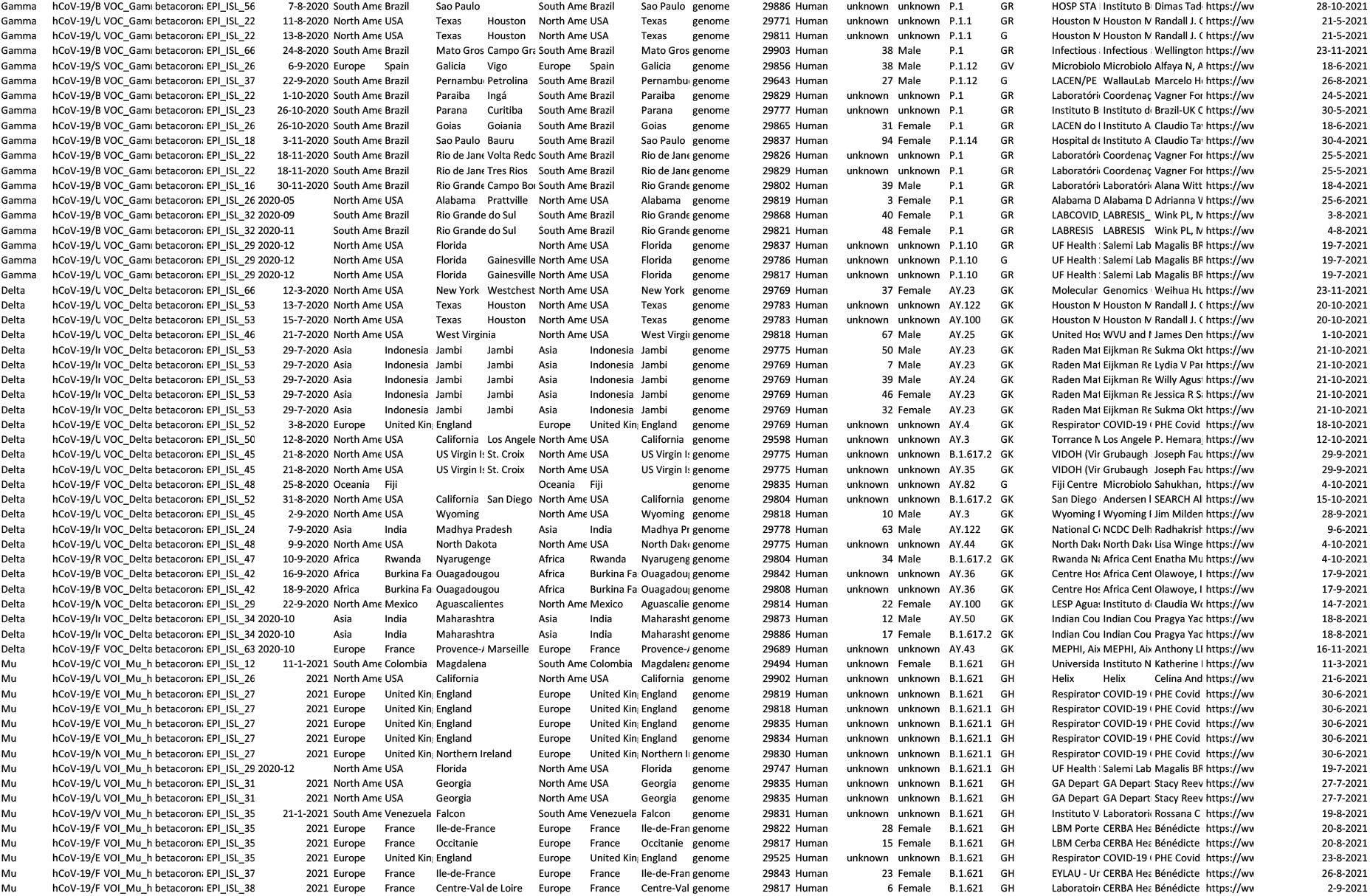
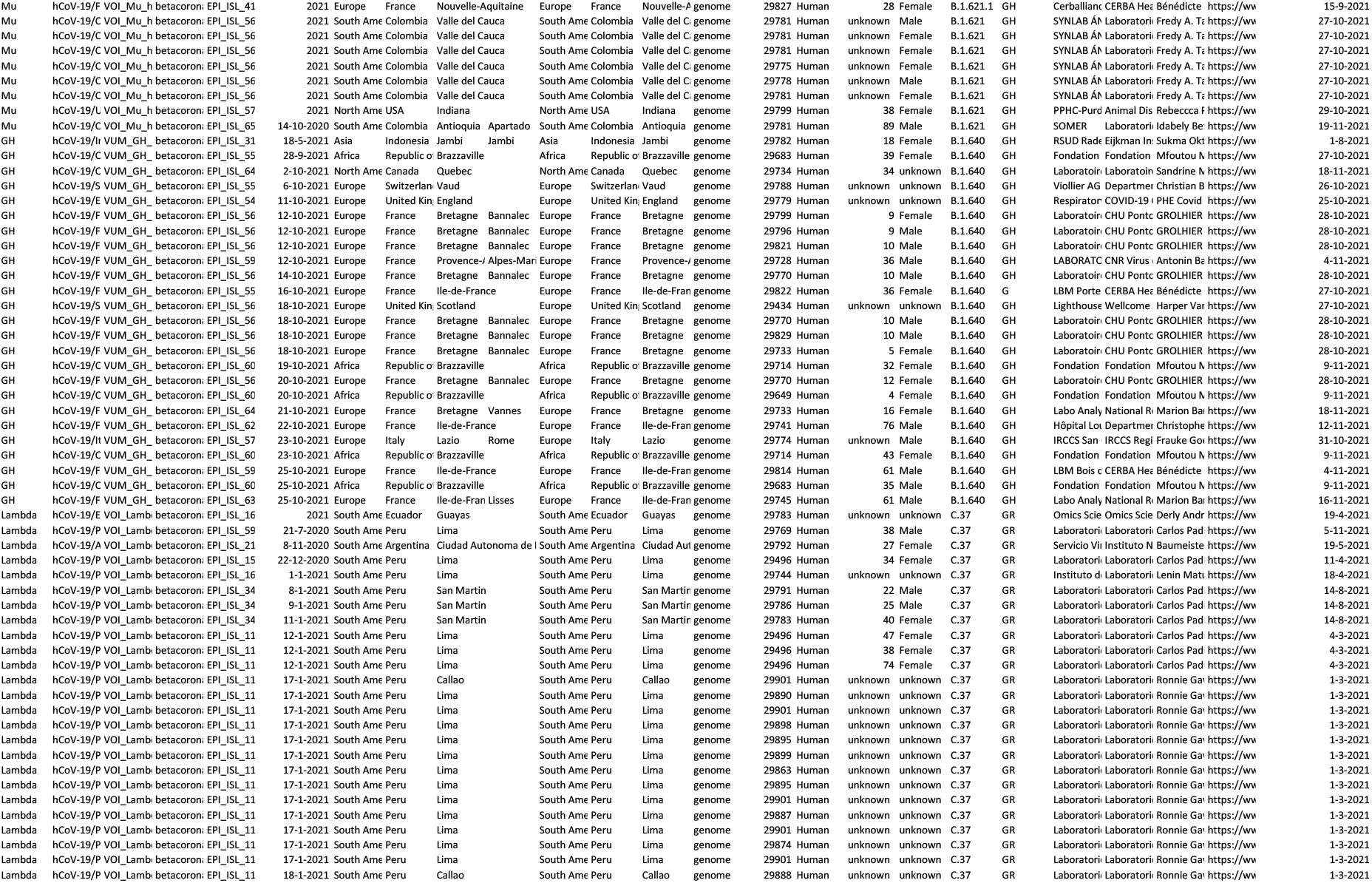
Metadata of the VOCs, VOIs and VUMs strains used in the present study.

**Table 2:**
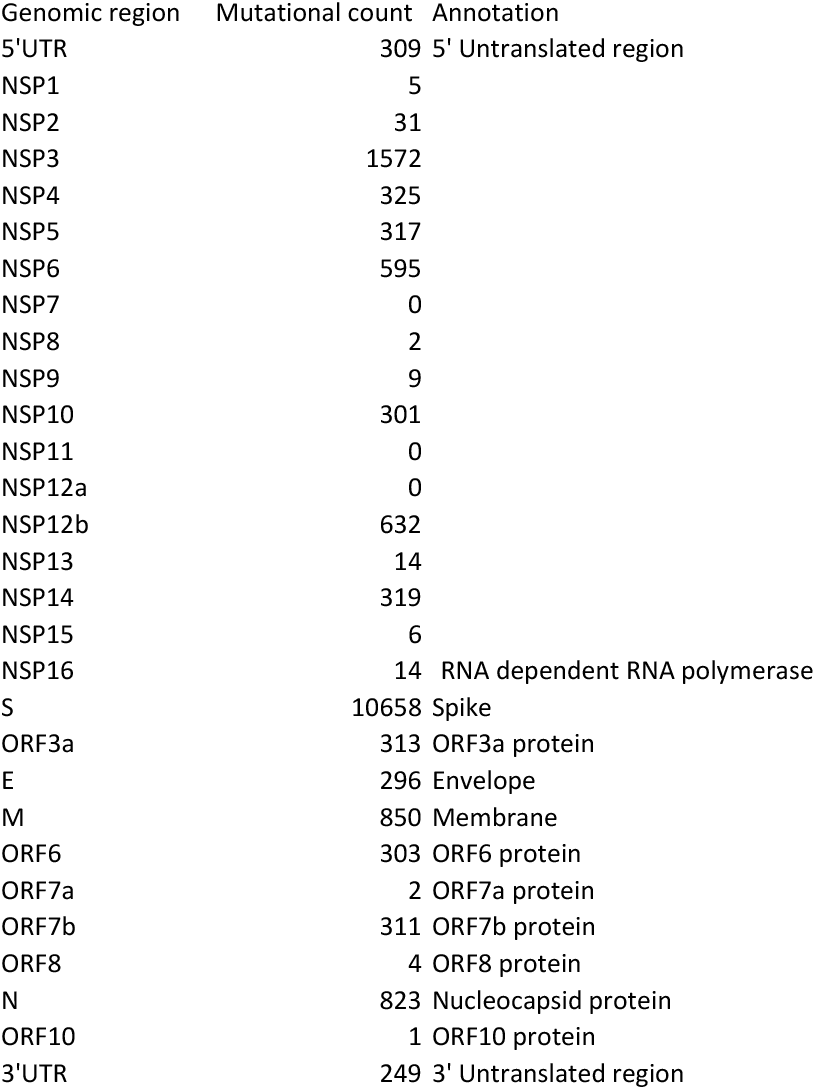
Genomic region wise mutational count of the omicron isolates by taking NC_045512.2 as a reference.

| Genomic region | Mutational count | Annotation |
| --- | --- | --- |
| 5'UTR | 309 | 5' Untranslated region |
| NSP1 | 5 |  |
| NSP2 | 31 |  |
| NSP3 | 1572 |  |
| NSP4 | 325 |  |
| NSP5 | 317 |  |
| NSP6 | 595 |  |
| NSP7 | 0 |  |
| NSP8 | 2 |  |
| NSP9 | 9 |  |
| NSP10 | 301 |  |
| NSP11 | 0 |  |
| NSP12a | 0 |  |
| NSP12b | 632 |  |
| NSP13 | 14 |  |
| NSP14 | 319 |  |
| NSP15 | 6 |  |
| NSP16 | 14 | RNA dependent RNA polymerase |
| S | 10658 | Spike |
| ORF3a | 313 | ORF3a protein |
| E | 296 | Envelope |
| M | 850 | Membrane |
| ORF6 | 303 | ORF6 protein |
| ORF7a | 2 | ORF7a protein |
| ORF7b | 311 | ORF7b protein |
| ORF8 | 4 | ORF8 protein |
| N | 823 | Nucleocapsid protein |
| ORF10 | 1 | ORF10 protein |
| 3'UTR | 249 | 3' Untranslated region |

**Table 3:**
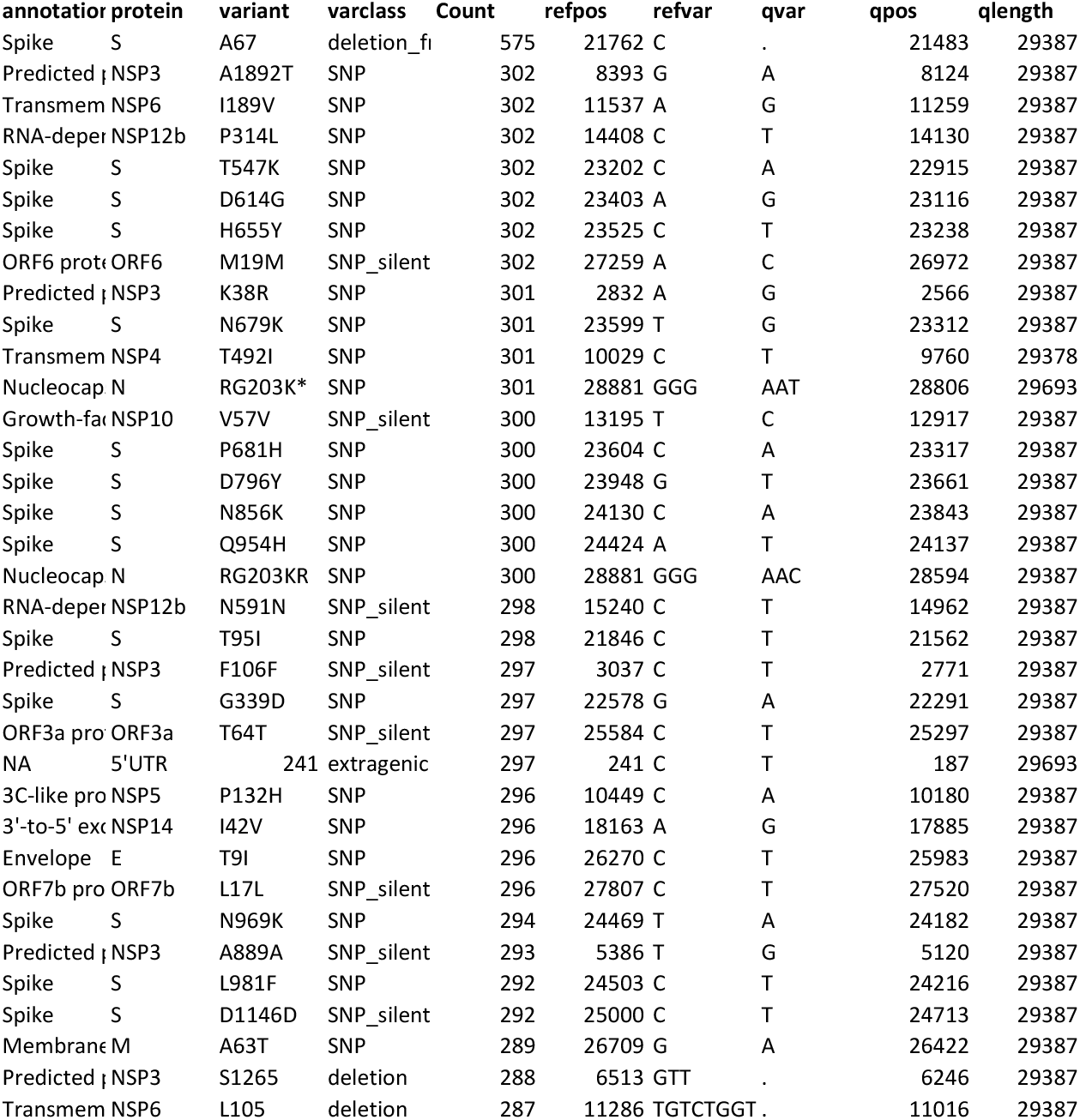
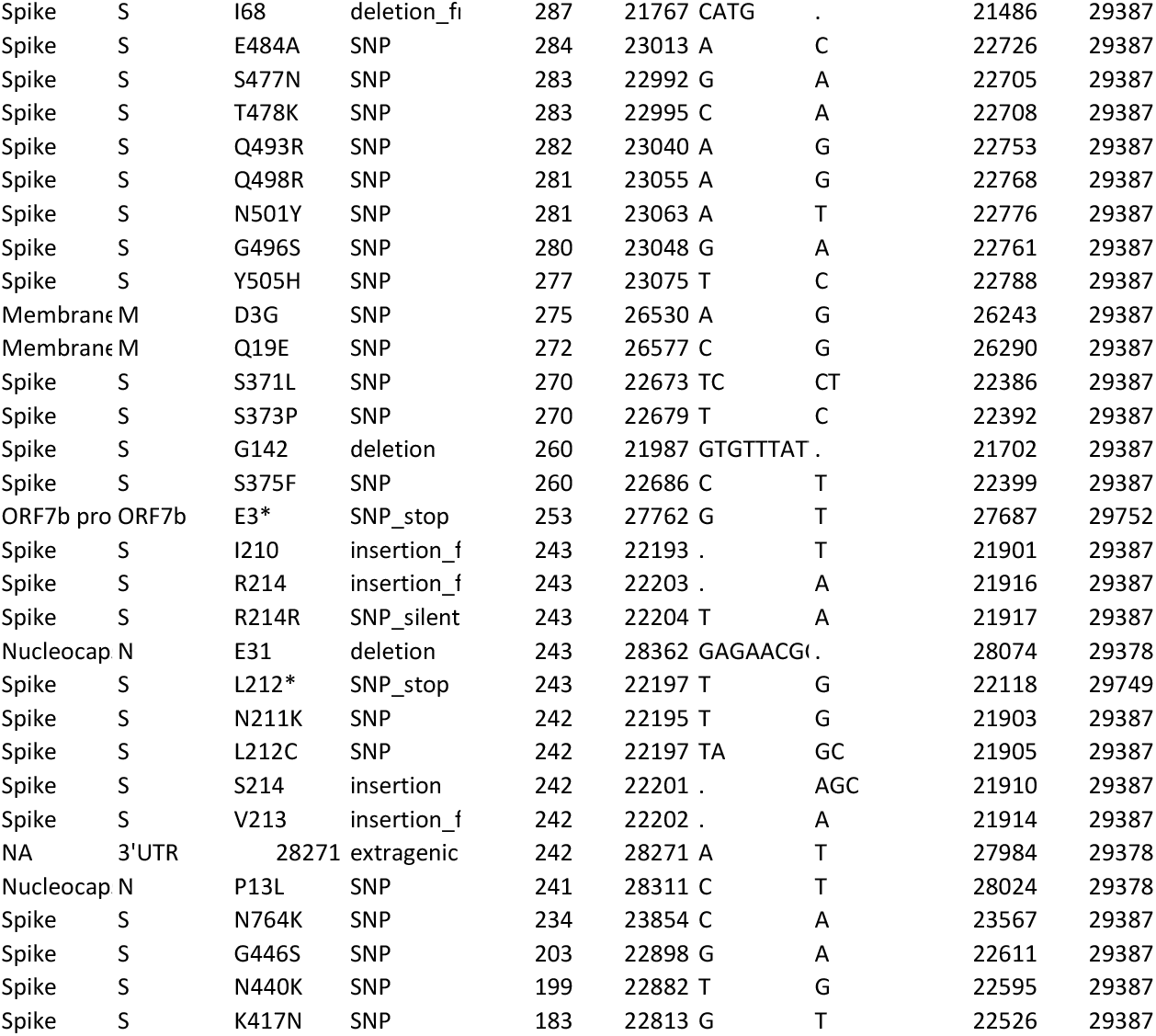
Top mutations (>185 in count) in omicron variant as compared to the reference sequence NC_045512.2

### Low intra-sequence diversity amongst omicron variant

Intra-strain diversity among the omicron variant strains reported worldwide will be crucial in understanding the genome dynamics and rapid evolution of SARS-JCoV-2. We performed the mutational analysis on the current dataset using omicron (OL677199) isolated from Canada on 23^rd^ November 2021 as the reference genome. Most of the strains (n = 298), irrespective of their geographic origin, had less than ten mutations depicting low intra-strain diversity among omicron strains. However, four of the isolates two from Europe (Italy) (EPI_ISL_6854347 (n = 23 mutations) and EPI_ISL_6854346 (n = 14 mutations) and two from South Africa (EPI_ISL_6699742 (n =12 mutations) and EPI_ISL_6774091 (n =11 mutations) were most diversified among the omicron genomes. While omicron had >55 mutations when compared with other VOCs and VOIs.

## Methods

### Identification and procurement of SARS-CoV-2 genome from the public repository

We have considered all the available genomes of omicron variant available in public domain until 6 pm Indian Standard Time (IST) on 2^nd^ December 2021 from GISAID (n = 302 genomes). A total of 25 strains from each variant of concern, namely alpha (B.1.1.7), beta (B.1.351), gamma (P.1) and delta (B.1.617.2) and variant of interest, namely lambda (C.37) and mu (B.1.621). We have also considered 25 strains from variant under monitoring, namely GH (B.1.640). These all strains are from their respective earlier reports in the public domain. A detailed list of all the strains used in the study is provided in the supplementary information (supplementary table 1).

### Phylogenetic analysis

A total of 477 high-quality genomes, including the major variants spread across the globe were taken into consideration. Multiple sequence alignment was performed for all the genomes using MAFFT v7.467 (Nakamura, Yamada, Tomii, & Katoh, 2018) followed by phylogenetic tree construction using fasttree v2.1.8 with double precision (Price, Dehal, & Arkin, 2010) with gamma time reversal method. Visualization of the obtained phylogenetic tree was performed using iTol v6 (Letunic & Bork, 2019). Different variants were marked in accordance with different colors as mentioned in the legends.

### Mutational analysis

Mutational analysis of all the strains (n=477) in the study was performed with two different reference genomes. First with NC_045512.2 (Wuhan-Hu-1) strain (reference SARS CoV-2 strain) and another with first reported strain of omicron variant (OL677199.1) (https://www.ncbi.nlm.nih.gov/nuccore/OL677199) using nucmer v3.1 (Delcher, Phillippy, Carlton, & Salzberg, 2002). We have used a well-documented R script described earlier (Mercatelli & Giorgi, 2020). Here, we have used gff3 annotation and reference genome file to extract genomic coordinate of SARS-CoV-2 proteins. R library package seqinr (https://cran.r-project.org/web/packages/seqinr/index.html) and biostring package (https://bioconductor.org/packages/release/bioc/html/Biostrings.html) of bioconductor was implemented to obtain the list of all the mutational events. Frequency and rate of mutation were calculated with respect to two different references (Reference SARS CoV-2 strain: NC_045512.2) (https://www.ncbi.nlm.nih.gov/nuccore/NC_045512.2) and omicron (OL677199.1) (https://www.ncbi.nlm.nih.gov/nuccore/OL677199) separately.

## Abbreviations

VOC: variant of concern
VOI: variant of interest
VUM: variant under monitoring
NSP: non-structural protein
UTR: untranslated region
rdrp: RNA dependent RNA polymerase

## Author contribution statement

Kanika Bansal performed data curation, analysis and writing of manuscript with inputs from Sanjeet Kumar.

## Acknowledgement

Authors acknowledge the support and motivation from Dr. Prabhu B.Patil – CSIR-Institute of Microbial Technology, Chandigarh. We are also thankful to Dr. Santosh Kumar Sethi for his kind support during the process of study. We also acknowledge GISAID initiative for extensive curation and availability of genomic resource in public domain.

## Supplementary material

**Supplementary figure 1: Maximum likelihood whole genome-based phylogeny of SARS-CoV-2 VOCs, VOIs and VUMs.** Here, phylogroups (PG-I and PG-II) and clades (alpha, beta, gamma, delta, omicron, mu etc.) are marked with respective colors as indicated. Bootstrap values are represented by the radius of circle at the nodes. For better visualisation of tree topology and bootstrap values branch length is ignored.

**Supplementary table 1:** Total mutations in strains under study using NC_045512.2 as reference genome.

**Supplementary table 2:** Total mutations in strains under study using omicron (OL677199) as reference genome.

