## Supplementary figures and images for "Mutational cascade of SARS-CoV-2 leading to evolution and emergence of omicron variant"

### supplementary figure 1

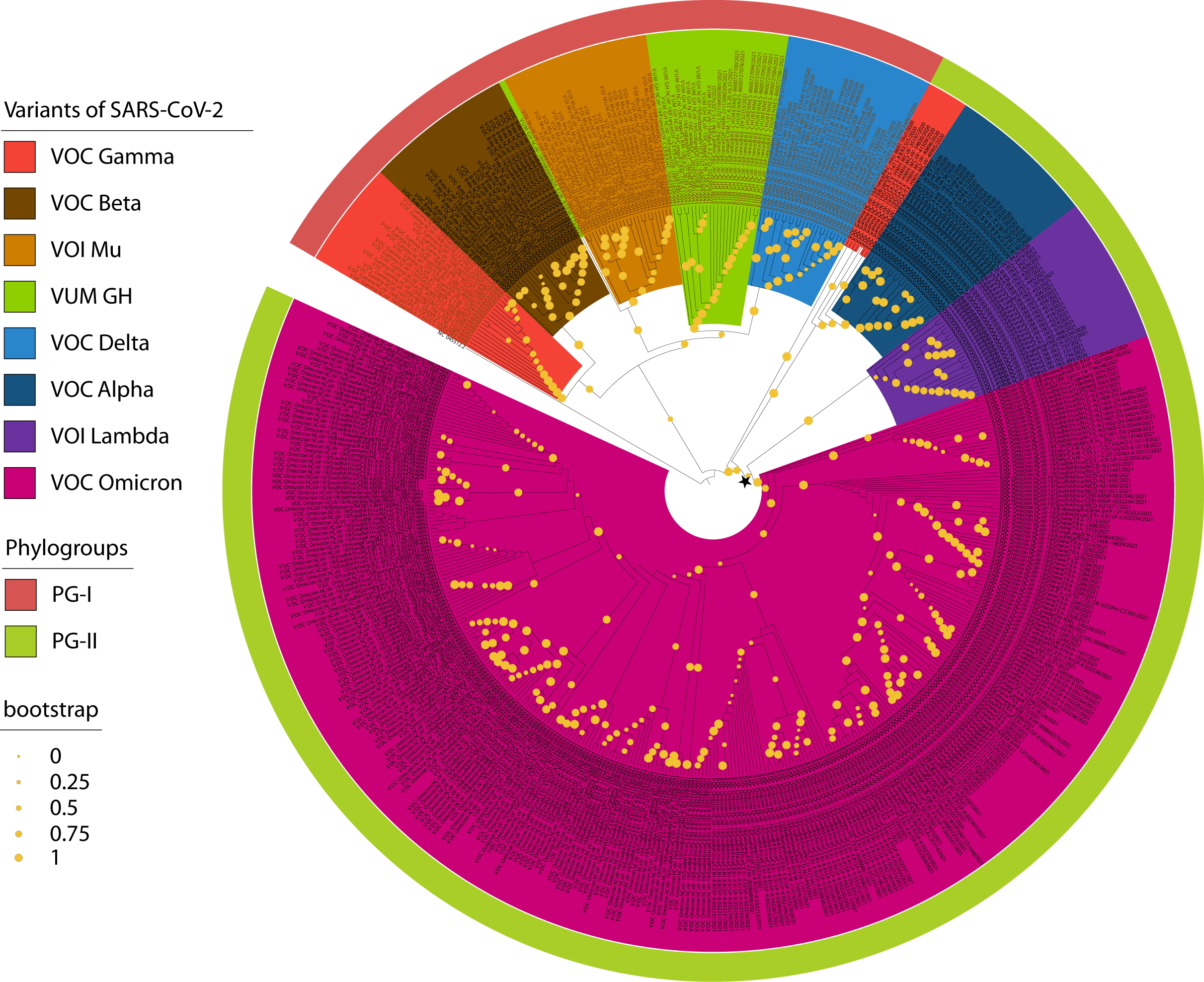
